# Ontogeny of superorganisms: Social control of queen specialization in ants

**DOI:** 10.1101/2022.03.08.483434

**Authors:** Vahideh Majidifar, Marina N. Psalti, Martin Coulm, Ebru Fetzer, Eva-Maria Teggers, Frederik Rotering, Judith Grünewald, Luca Mannella, Maxi Reuter, Dennis Unte, Romain Libbrecht

## Abstract

A central question in life sciences is to understand the ontogeny of biological systems, which exist at multiple phenotypic scales and function via the cooperation of specialized entities. Examples of such systems include multicellular organisms, which consist of specialized cells, and insect societies (or superorganisms), which are composed of specialized individuals. Both systems are products of major evolutionary transitions, and they share commonalities in their ontogeny, as both develop from a single, pluripotent unit. While the ontogeny of multicellular organisms is well understood, the factors and mechanisms that control the ontogeny of superorganisms remain poorly studied. Here, we report experimental investigations of the process of colony foundation in ants. In most ant species, a new colony is established by a solitary founding queen that expresses behavioral pluripotency to produce the first workers, at which point the queen becomes strictly specialized in egg production. We demonstrate that the presence of workers is necessary and sufficient to induce this specialization of queens. Moreover, workers also maintain the queen specialization in mature colonies, as established queens isolated from their workers revert to expressing behavioral pluripotency. Our results also suggest that this underappreciated social control of queen specialization may be common in ants and regulated by ancestral mechanisms. These findings stand in contrast to the traditional view of social insect queens as being intrinsically specialized in egg production and may reshape our understanding of division of labor in insect societies.

**Significance statement:** Insect societies are characterized by division of labor between queens that specialize in producing eggs and workers that perform all non-reproductive tasks. Studies of division of labor traditionally focused on fully established colonies and there is limited information on the factors and mechanisms that initiate division of labor during colony foundation. Here, we report that the presence of workers not only initiates the queen specialization, but also maintains it continually throughout the colony life. Finding such a social control of the specialization of queens contradicts the commonly accepted view of social insect queens as intrinsically specialized egg-laying machines. Our study has the potential to reshape our understanding of the functioning and evolution of insect societies.

---

The functioning of biological systems relies on the cooperation of specialized components and understanding the emergence and evolution of this specialization is a major challenge in biology. Typical examples of such biological systems include multicellular organisms that are composed of specialized cells, and insect societies that are composed of specialized individuals(1). Indeed, social insect colonies (also called superorganisms) are analogous to multicellular organisms in that they have queens that monopolize reproduction (similar to germ cells), and functionally sterile workers that perform all non-reproductive tasks and thus act as somatic cells (2, 3). Both types of biological systems evolved from solitary and non-specialized ancestors in major evolutionary transitions (1): multicellular organisms from unicellular organisms, and superorganisms from solitary insects.

Interestingly, in both cases, the specialization also needs to be established in every generation during the ontogeny of these biological systems (i.e., the developmental process that produces the self-assembly and specialization of their components from a single unit). Extensive research efforts have focused on the ontogeny of multicellular organisms, building up the entire field of developmental biology. This demonstrated that studying the ontogeny is a powerful approach to understand the evolution and emergence of specialization (4, 5). However, this approach has not been applied to social insects, and there are few experimental investigations of the ontogeny of insect societies (6, 7).

In most social insect species, mated queens found their colony independently (8), and are similar to zygotes in that they are the earliest developmental stage of superorganisms (2). The development of colonies from founding queens is the ontogeny of superorganisms. These pluripotent founding queens express a broad repertoire of behaviors and fulfill multiple functions to produce the first workers (6, 9–14). It is only once the colonies are established (i.e., they contain workers) that the queens become strictly specialized in egg production (15). While the queen specialization is a central process in the ontogeny of superorganisms, as is cell differentiation in the ontogeny of multicellular organisms, the factors and mechanisms that control the specialization of pluripotent founding queens remain poorly understood.

In this study, we investigate queen specialization in terms of brood care behavior. While workers provide brood care in mature colonies, queens need to do so during the founding stage because the brood of social Hymenoptera cannot develop independently (16). Using the black garden ant *Lasius niger* as a study system, we show that the presence of workers is necessary and sufficient to inhibit brood care in queens, and thus initiate their behavioral specialization. We find this specialization to be reversible, as queens revert to expressing brood care upon experimental removal of their workers. Additional experiments suggest that such behavioral flexibility of queens may be conserved across ants. By revealing a continual social control of queen specialization, our results jeopardize the common conception of ant queens as being intrinsically specialized egg-laying machines.

## Results and Discussion

### Queens become specialized after the emergence of the first workers

To verify that *L. niger* founding queens raise their first cohort of workers independently, we collected newly mated queens right after their nuptial flight and transferred them to closed, individual glass tubes. We monitored brood production and development over the next 93 days. 91.67% of the queens (22/24) survived the experiment, and 77.28% of the surviving queens (17/22) produced workers within the observation time (5.26±3.95 workers; mean±sd; Fig. S1). This indicates that founding queens provide care to the brood, as ant larvae cannot develop independently (16). We confirmed this by direct observations of brood care behavior (defined as active manipulation of the brood) in founding queens before and after worker emergence. We found that founding queens expressed brood care before they produce the first workers, but that the emergence of workers was associated with a sharp decrease in queen brood care behavior (χ^2^=35.63, P<0.0001, Fig. 1A). Therefore, the emergence of workers correlates with the behavioral specialization of queens in *L. niger*.

**Fig. 1.**
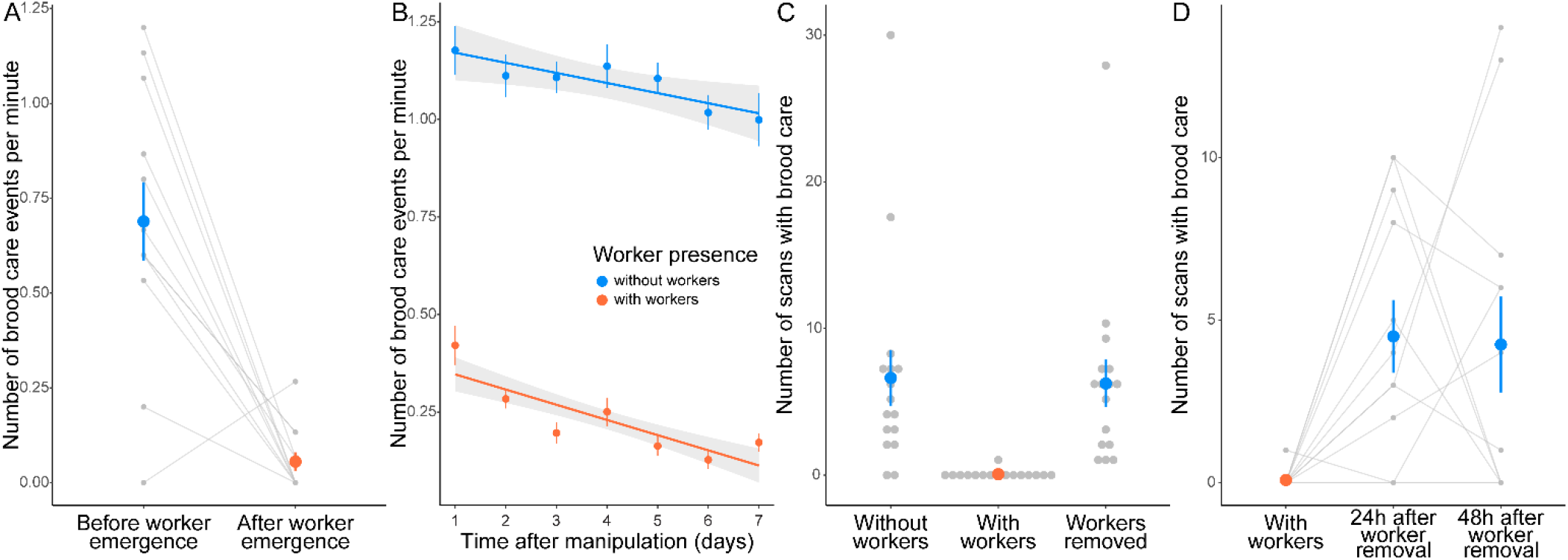
The presence of workers both initiates and maintains the queen behavioral specialization. (**A**) The emergence of workers correlates with the behavioral specialization of queens. Founding queens (n=12) reduced brood care after producing the first workers (χ^2^=35.63, P<0.0001). (**B**) The experimental addition of workers drives the queen specialization. Founding queens provided with workers (n=29) showed lower brood care levels than founding queens kept without workers (n=25, χ^2^=596.33, P<0.0001). The effect of worker presence was already detected 20 hours after the addition of workers (χ ^2^=221.28, P<0.0001). (**C**) The queen specialization is reversible after three days. Founding queens that had their workers removed (n=16) performed more brood care than queens that just received workers (n=16, t=6.19, P<0.0001), but similar brood care as queens that never had any workers (n=16, t=0.15, P=0.99). (**D**) The queen specialization is reversible after two years and six months. Established queens (n=12) showed increased brood care both 24 hours (t=3.69, P=0.0035) and 48 hours (t=3.07, P=0.015) after their workers were removed. Colored dots and error bars represent means and standard errors, respectively, and grey dots show individual data points.

### The presence of workers triggers the specialization of queens

The emergence of workers is confounded with the age and nutrition of queens, as established queens - defined as queens with workers - are older and fed by their workers. To disentangle age and worker presence, we developed a protocol to provide workers to same-age founding queens. We sampled brood from field colonies and kept it in laboratory colonies that we monitored regularly to collect workers that recently emerged from the pupae. Such workers are readily accepted by foreign ants (6, 17–20), possibly because they do not possess the signature chemical profile of their own colony yet (21–23). We manipulated worker presence and feeding status of same-age founding queens that had not produced workers yet, and quantified their brood care behavior for seven days. We detected an interaction between time and worker presence (χ^2^=17.24, P<0.0001), as queens with workers showed a stronger decrease in brood care over time (χ^2^=49.69, P<0.0001, Fig. 1B) than queens without workers (χ^2^=8.99, P=0.0027, Fig. 1B). In addition, we found very strong evidence that queens with workers performed less brood care overall than queens without workers (χ^2^=596.33, P<0.0001, Fig. 1B). This inhibition of brood care by workers was already detected 20 hours after the addition of workers (χ^2^=221.28, P<0.0001, Fig. S2). We could not detect any effect of the feeding status, neither as a main effect (χ^2^=1.10, P=0.29) nor as an interaction with time (χ^2^=1.90, P=0.17). This experiment demonstrates that the presence of workers is necessary and sufficient to initiate the behavioral specialization of *L. niger* queens.

### The presence of workers maintains the specialization of queens in established colonies

Then, we investigated whether the presence of workers is necessary to maintain the queen specialization in established colonies. First, we studied this question in *L. niger* queens that were provided with workers for three days. As expected, those queens expressed lower levels of brood care than same-age queens that never had any workers (χ^2^=12.78, P=0.00035). We then removed the workers and compared the queen brood care behavior to queens that either never had any workers or were just provided with workers. We found that queens that had their workers removed performed more brood care than queens that just received workers (t=6.04, P<0.0001, Fig. 1C), but similar levels of brood care as queens that never had any workers (t=0.15, P=0.99, Fig. 1C).

We report that the worker-driven inhibition of queen brood care behavior is reversible after three days, but it may be that more time is needed for the queen specialization to be permanently established. Therefore, we investigated whether the queen behavioral specialization is reversible after several years. To do so, we used *L. niger* colonies that were founded in the laboratory two years and six months before starting the experiment. We first recorded the queen brood care behavior after removing all but five workers. Then, we removed the last five workers, and quantified queen brood care behavior on the next two days. We found that queens expressed elevated brood care levels both 24 hours (t=3.69, P=0.0035, Fig. 1D) and 48 hours (t=3.073, P=0.015, Fig. 1D) after worker removal. We did not detect any difference between the two time points (t=0.61, P=0.81, Fig. 1D), indicating that the queens did not show further behavioral changes after 24 hours. To complete these findings, we quantified changes in queen brood care behavior in response to the sequential removal of workers in laboratory colonies that were established three years and two months before the experiment. We found that these established queens went back to expressing brood care upon the experimental removal of their last worker (Fig. S3). These experiment show that the presence of workers not only initiates the specialization of *L. niger* queens during colony foundation, but also constantly maintains it in established colonies.

### Mating is not required for queens to become specialized in response to workers

Then, we investigated whether mating and reproductive activity are pre-requisites for queens to express brood care behavior, and whether it depends on worker presence. To do so, we manipulated the presence of workers and quantified the brood care behavior of winged unmated queens collected from their colonies of origin (thus before their nuptial flight). We found that unmated queens performed overall very low levels of brood care behavior (Fig. S4). Such expression of brood care by unmated queens has been reported in other ant species (24–26). In addition, we found moderate evidence that the likelihood of expressing brood care was lower for unmated queens kept with workers compared to unmated queens kept without workers (χ^2^=4.50, P=0.034, Fig. S4). These results indicate that the presence of workers triggers the specialization of queens independent of their mating status and reproductive activity.

### Workers induce queen specialization in a dose-dependent manner

After demonstrating that the presence of workers controls the queen specialization, we set out to characterize the inhibitory effect of workers on brood care. First, we investigated whether the effect of workers is dose-dependent, as in other instances of social control of phenotypic variation in social insects (27, 28). We provided same-age founding queens with five larvae, and either zero, one, two, three or five workers, and found a negative, non-linear correlation between queen brood care and the number of workers (t=8.45, P<0.0001, Fig. 2A). While the presence of a single worker was sufficient to drive a reduction in brood care behavior, additional workers inhibited brood care further, as the correlation remained after removing the queens that did not receive any workers from the analysis (t=3.35, P=0.0014). We repeated the same experiment with 15 larvae instead of five - thus increasing the larval need for care - and also found a negative correlation between the number of workers and the level of brood care expressed by queens, both when queens without workers were included (t=4.19, P<0.0001, Fig. S5) and excluded from the analysis (t=2.37, P=0.021). These results indicate that workers have a dose-dependent negative effect on the brood care behavior of queens.

**Fig. 2.**
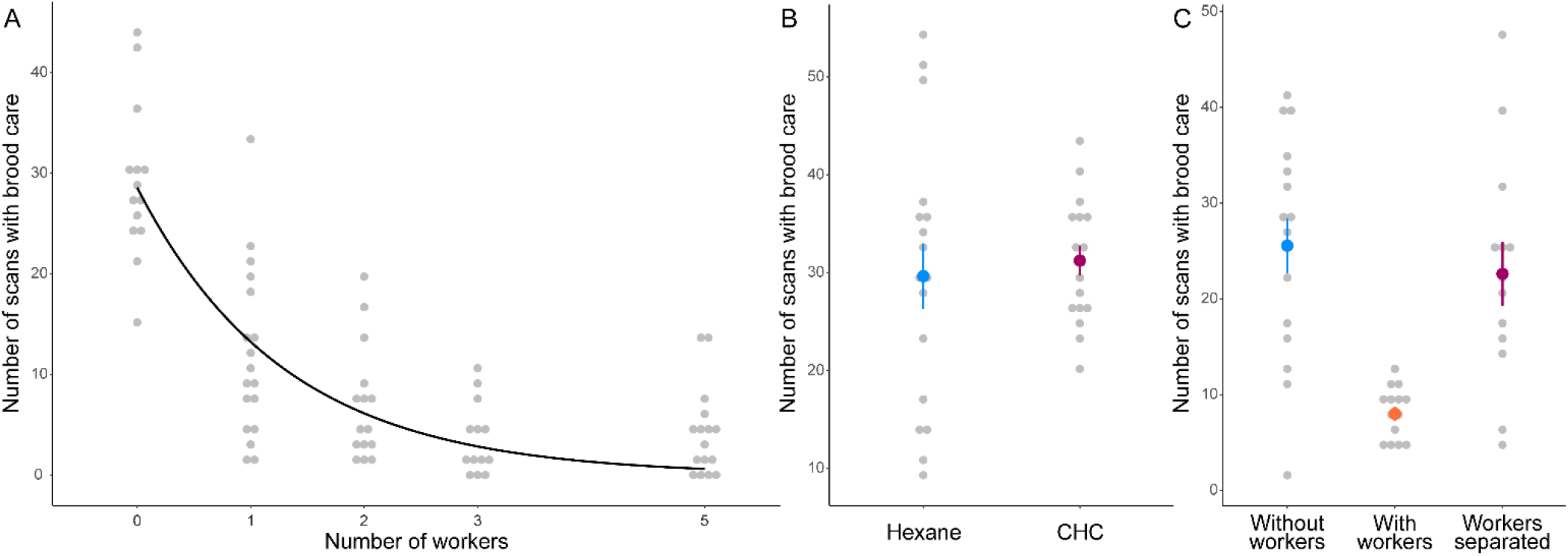
(**A**) Workers show a dose-dependent inhibition of queen brood care. Founding queens were provided with zero (n=14), one (n=18), two (n=15), three (n=13) or five (n=16) workers. There was a negative, non-linear correlation between the number of workers and the queen brood care level (t=8.5, P<0.0001). The black line shows the non-linear regression curve of the model, and grey dots show individual data points. (**B**) Worker cuticular hydrocarbons (CHC) do not drive the queen specialization. No behavioral difference was detected depending on whether founding queens received glass beads covered in worker CHC (n=18) or hexane (control, n=18, χ^2^=0.60, P=0.44). (**C**) Workers separated by a wire mesh do not drive the queen specialization. The brood care behavior of founding queens kept with workers separated by a wire mesh (n=21) differed from founding queens kept with workers (n=14, t=3.95, P=0.001), but not from founding queens kept without workers (n=20, t=0.66, P=0.79). Colored dots and error bars represent means and standard errors, respectively, and grey dots show individual data points.

### Mere worker cues do not induce the specialization of queens

Dose-dependent effects of social partners on phenotypic variation are often driven by variation in the quantity of social cues. Because social insects detect social partners via the blend of hydrocarbons on their cuticle (29, 30), we extracted cuticular hydrocarbons (CHC) from pools of *L. niger* workers and applied the CHC to glass beads that we provided to same-age founding queens. We did not detect any effect of the CHC treatment on the expression of brood care by queens (χ ^2^=0.60, P=0.44, Fig. 2B), indicating that queens do not modify their behavior in response to the mere detection of worker CHC. This result was confirmed by three additional experiments. First, we did not detect any difference in brood care behavior between founding queens depending on whether they were kept in boxes that used to contain many workers, or in boxes that never contained any (χ^2^=0.48, P=0.49, Fig. S6). Second, we found that the presence of frozen workers, and thus of worker CHC (31–33), had no detectable effect on queen brood care (t=0.57, P=0.84, Fig. S7). Finally, we investigated how the queen behavior was affected by workers separated from the queen and brood by a wire mesh. This setup enabled workers to make antennal contacts with the queen and brood, but prevented closer interactions such as fluid exchange via trophallaxis (34). We found that queens kept with workers separated by a wire mesh expressed more brood care than queens kept together with workers (t=3.95, P=0.001, Fig. 2C), but as much brood care as queens kept without workers (t=0.66, P=0.79, Fig. 2C). This series of experiments shows that worker cues are not sufficient to drive the queen specialization and suggests that workers require close interactions with queens and/or larvae to inhibit the brood care behavior of queens.

### Juvenile hormone may regulate the specialization of queens

The juvenile hormone (JH) pathway is a good candidate for the regulation of brood care in queens because it plays such a role in workers of some social insect species (35, 36). To investigate the role of JH, we observed the behavioral response of same-age founding queens (kept with or without workers) to treatments with a JH analog (methoprene) or with chemical inhibitors of JH production (precocene I and II). We found that the methoprene treatment decreased queen brood care in the first two days of the experiment, when brood care levels were relatively high (χ^2^=4.43, P=0.035, Fig. 3A), consistent with previous reports of methoprene inhibiting brood care in *L. niger* queens (36). On the contrary, we found that queens treated with the JH inhibitor precocene I expressed elevated levels of brood care, although this was only detectable on the third and fourth day of the experiment, when brood care levels were overall low (χ^2^=3.91, P=0.048, Fig. 3B; see Fig. S8 and Table S1 for details on the effect of precocene II). These effects of methoprene and precocene I were independent of worker presence (Table S1). Finding that treatments with a JH analog reduced brood care, while treatments with a JH inhibitor increased brood care, suggests a possible role of the JH pathway in regulating the brood care behavior of *L. niger* queens.

**Fig. 3.**
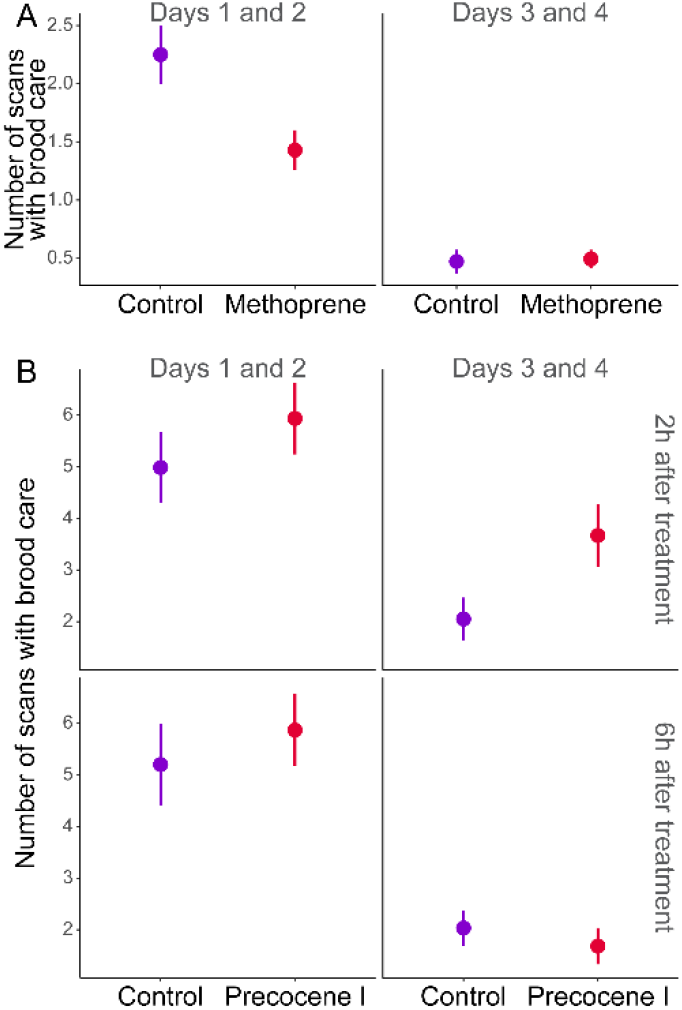
A possible role of JH in regulating the brood care behavior of queens. (**A**) Founding queens treated with methoprene (JH analog, n=35) showed reduced brood care levels compared to control queens (acetone, n=35) on days 1 and 2 (χ ^2^=4.43, P=0.035), but not on days 3 and 4 (χ^2^=1.05, P=0.31). (**B**) Queens treated with precocene I (JH inhibitor, n=30) increased brood care levels compared to control queens (acetone, n=30) on days 3 and 4 (χ^2^=3.91, P=0.048), but not on days 1 and 2 (χ^2^=2.34, P=0.13). The effect on days 3 and 4 was primarily driven by the difference between treated and control queens two hours after treatment (χ^2^=10.49, P=0.0012), which was not detectable anymore six hours after treatment (χ^2^=0.41, P=0.52).

### Founding queens show an ancestral physiological response to the presence of larvae

The JH pathway interacts closely with the insulin-signaling pathway (37–40), and both show variation between queens and workers in various social insect species (38, 41–44). In ants, the insulin-signaling pathway likely played a role in the emergence of reproductive division of labor (45) from a subsocial ancestor that would alternate between reproduction and brood care phases (46–49). These phases of the ancestral life cycle were controlled by the brood, which inhibited reproduction (50). To investigate the effect of brood on the reproduction of founding and established queens, we quantified the effect of larvae on egg production over time in isolated *L. niger* queens before and after they produced workers. We found that, while the presence of larvae stimulated egg production in established queens (interaction between time and presence of larvae: χ^2^=50.02, P<0.0001; main effect of the presence of larvae: χ^2^=9.76, P=0.0018; Fig. 4A), it had the reverse effect and inhibited egg production in founding queens (interaction between time and presence of larvae: χ^2^=68.84, P<0.0001; main effect of the presence of larvae: χ^2^=16.47, P<0.0001; Fig. 4A). Because larvae inhibit their egg production, founding queens show an ancestral physiological response to the presence of brood, and are thus more similar in that respect to the subsocial ancestor of ants than to established queens.

**Fig. 4.**
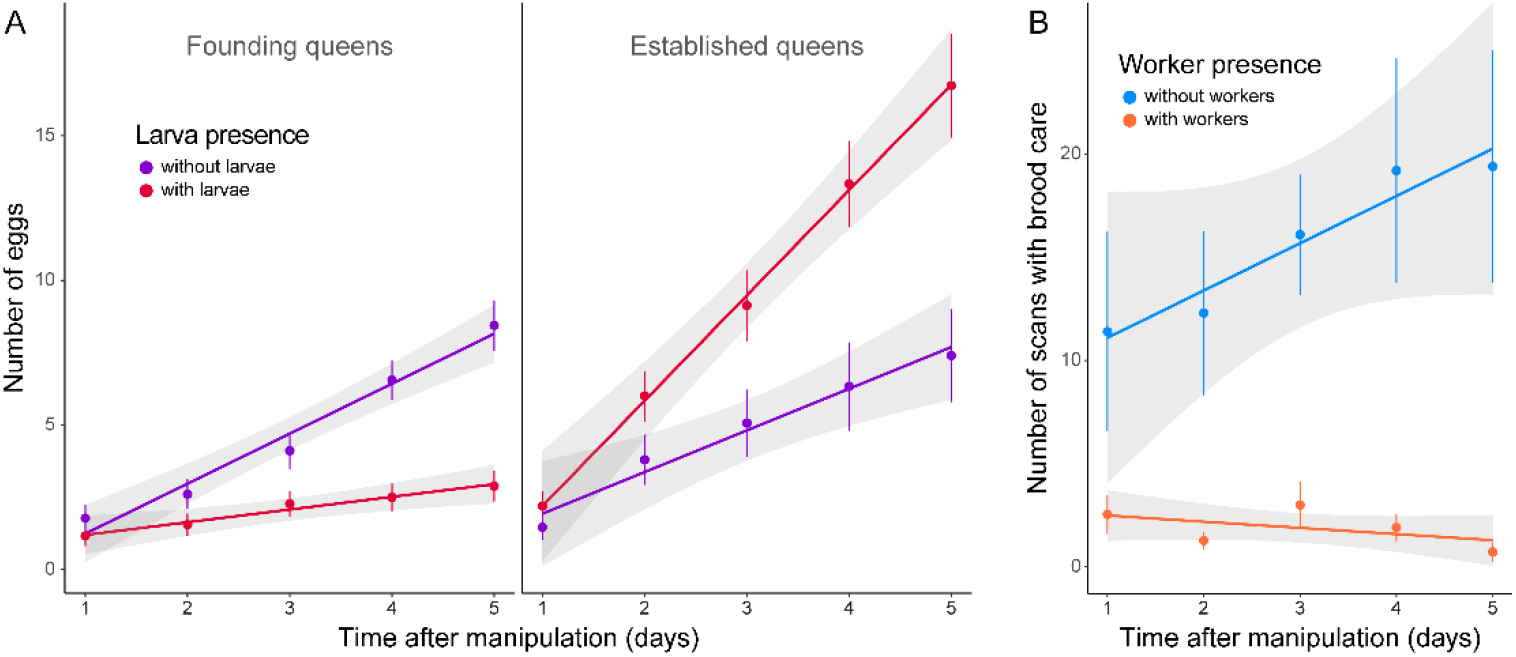
(**A**) Founding queens express an ancestral physiological response to brood. Founding queens (left) kept with larvae (n=18) produced less eggs than founding queens kept without larvae (n=18; interaction between time and presence of larvae: χ^2^=68.84, P<0.0001; main effect of the presence of larvae: χ^2^=16.47, P<0.0001), while established queens (right) kept with larvae (n=15) produced more eggs than established queens kept without larvae (n=15; interaction between time and presence of larvae: χ^2^=50.02, P<0.0001; main effect of the presence of larvae: χ^2^=9.76, P=0.0018). Note that both founding and established queens were kept without workers for the duration of the experiment. (**B**) Workers drive the queen specialization in another ant species. Established queens of the ant *Temnothorax nylanderi* expressed more brood care behavior when kept without (n=10) than with workers (n=11; χ^2^=32.23, P<0.0001). Colored dots and error bars represent means and standard errors, respectively.

### Workers induce the queen specialization in multiple species of ants

After finding that founding queens may express ancestral traits, we hypothesized that the social control of queen specialization may not be specific to *L. niger*, but rather occurs across the phylogeny of ants. To test this, we used field-collected mature colonies of *Temnothorax nylanderi* (which belongs to a different subfamily from *L. niger*), and experimentally manipulated the presence of workers. In addition to worker presence affecting queen brood care changes over time (interaction between time and worker presence: χ^2^=3.96, P=0.047), we found that queens without workers generally expressed more brood care than queens with workers (χ^2^=32.23, P<0.0001, Fig. 4B). This result shows that workers maintain queen specialization in *T. nylanderi* as well, and together with previous reports of social effects on queen behavior (6, 9, 51), suggests that it may be a common feature in social insects (but see (10, 52)).

## Conclusion

Our study reveals several commonalities between the ontogeny of superorganisms and multicellular organisms. First, we demonstrate and characterize the central role of the social environment in initiating and maintaining queen specialization, which mirrors the importance of cell-cell interactions in the process of stem cell differentiation in multicellular organisms (53–58). Second, we found that the specialization of ant queens is reversible, as queens kept without workers revert to expressing behavioral pluripotency. While this finding seems to contrast with the traditional view of stem cell differentiation as being irreversible, it is aligned with the accumulating evidence of experimental and natural dedifferentiation of specialized cells in some multicellular organisms (59–61). Third, we find that the physiological response to brood is similar between founding queens and the subsocial ancestor of ants, which echoes recent reports of molecular and structural similarities between pluripotent stem cells in multicellular organisms and their unicellular ancestors (4, 5, 62, 63).

Both insect societies and multicellular organisms evolved from pluripotent ancestors that had the ability to fulfill different functions (5, 46). Major evolutionary transitions resulted in these functions to be carried out by different, specialized entities that cooperate to operate at a higher phenotypic scale (1). Thus, despite specificities of superorganismic systems compared to multicellular organisms (e.g., the relatively weaker physical adhesion of their components), our findings draw interesting parallels between biological systems that exist at different phenotypic levels and originated from distinct major evolutionary transitions. This shows the importance of studying the ontogeny of superorganisms to expand our understanding of the ontogeny of biological systems beyond multicellular organisms.

Several lines of evidence suggest that the behavioral flexibility and physiological response to brood of the subsocial ancestor of ants are still apparent during colony foundation. First, the diverse behavioral repertoire of founding queens, as well as the effect of workers on queen behavior, can be found in multiple species of ants (9, 12, 16, 64), although not in all species that were investigated (10). Second, larvae inhibit egg production in founding queens, as it likely did in the ancestor of ants (50). Not having access to this subsocial ancestor has hampered the study of the evolution of eusociality in ants (65) and investigating pluripotent founding queens may provide a window into these ancestral functions. In that perspective, the transition from pluripotency to specialization that underlies colony foundation may even mirror the evolutionary transition from subsocial pluripotency to division of labor between specialized castes.

Finally, we report that queen specialization is reversible in two ant species that diverged more than 100 million years ago (66). This finding is unexpected because queens are typically seen as the fixed, intrinsically specialized caste that focuses solely on egg production (67). In addition, both *L. niger* and *T. nylanderi* are derived ant species with high degrees of social complexity and differentiation between the queen and worker castes. We propose that, although workers continuously maintaining queen specialization has remained relatively unnoticed so far, it may actually be a common feature of ant colonies, and possibly of other insect societies (6, 9, 51). By revealing such an underappreciated feature of division of labor and behavioral specialization, this study could reshape our understanding of the evolution and functioning of insect societies.

## Methods

### General procedures

Below we describe the general procedures regarding the collection and keeping conditions of queens, the use of callow workers for experimental manipulation of worker presence, the experimental setup and behavioral observations, as well as the statistical analyses.

#### Founding queen collection and keeping conditions

All *Lasius niger* founding queens used in the experiments were collected after their nuptial flights in 2017, 2018, 2019 and 2020 around Mainz and Ingelheim, Germany (Supplementary Table 2). After collecting them in fluon-coated boxes containing humid paper towel, we transferred the founding queens individually into glass tubes (10cm length x 1cm diameter) half filled with water blocked by cotton, and plugged with another piece of cotton. The tubes were placed in boxes (18cm x 12cm x 7cm) kept in darkness. Most queens were kept in a climate chamber at 21°C and 80% humidity. Founding queens were not provided with food, since *L. niger* queens found their colonies without foraging (claustral colony founding) (8, 68, 69). Once the first workers emerged, each tube containing the queen, brood and workers was transferred into a small plastic box (15cm x 11cm x 12cm) with walls coated with fluon to prevent the ants from escaping. These colonies were also kept in a climate chamber at 21°C and 80% humidity, but fed every second week with a drop of honey, one frozen cricket and a piece of artificial food with a 2:1 (carbohydrate:protein) ratio (70). Every winter, the established colonies were hibernated for three to four months at 5°C. They entered and left hibernation via a two-week gradual decrease and increase of temperature, respectively. Supplementary Table 2 provides information on queen collection, keeping conditions, and sample sizes in all experiments.

#### Experimental manipulations of worker presence

To experimentally provide workers to founding queens, we used “callow” workers (6, 17–20, 71, 72). Callow workers had emerged from the pupae very recently (< 12h), did not bear their colony-specific odor yet (21–23), and thus did not elicit aggression from foreign individuals (including founding queens). To obtain callow workers, we sampled brood from field colonies in and around of Mainz, kept it in laboratory colonies that we monitored regularly to collect recently emerged workers. Callow workers are easily recognizable due to their light grey color. The laboratory colonies used for callow worker production were kept between 21°C and 28°C, and between 80% and 100% humidity, depending on timing and availability of climate cabinets. In cases where callow workers were kept at a different temperature than the temperature of the experiment, they were moved to the room of the experiment at least one hour before it started.

#### Experimental setup

All behavioral analyses were conducted based on observations of videos that were recorded for one hour with cameras standing ca. 50cm over the observation arenas. Two white LED light bars illuminated the arenas approximately 70cm above the observation arenas. One camera recorded one tray, which could contain up to 12 observation arenas simultaneously (Supplementary Figure 9). Whenever multiple cameras were used simultaneously, the recordings belonged to the same batch. All experiments were filmed in a climate chamber at 21°C and 80% humidity, with 12h/12h light/dark cycle. The observation arenas consisted of airtight petri dishes (50mm diameter, 9mm height; Falcon) half filled with moistened, blue plaster for better contrast and visibility (Supplementary Figure 10). We used soft forceps and brushes to manipulate the ants. We ensure that all treatments of an experiment were represented and equally distributed on each tray. We assigned random numbers to the observation arenas to ensure that the experimenters were blind to the treatment during the experiments and video analyses.

#### Behavioral analyses

To assess the behavioral specialization of queens, we observed their brood care behavior toward larvae. We defined a brood care event as an active manipulation of the brood, with frequent contacts with the antennae and touches with the front legs. We also counted as a brood care event the placement of a brood item into a different position. However, merely carrying the brood item in the mouth parts was not scored as brood care. In the experiments “Worker emergence” and “Worker presence and feeding status”, we counted the total number of brood care events and divided it by the duration of the videos. In all other experiments, we scored the brood care behavior via scan sampling. We watched the first ten seconds of every minute of each video, and recorded the presence or absence of brood care behavior. We performed 51 such scans per 1h video, as we did not analyze the first and last five minutes of the videos to avoid potential disturbances due to the experimental manipulations. The videos were analyzed using the softwares BORIS (version 8.0.5) or Pot player (version 1.7.21212).

#### Statistical analyses

We performed all statistical analyses using R v. 4.0.4 (R Core Team 2021) and RStudio v. 1.4.1106 (R Core Team 2021). Supplementary Table 1 provides the input data, syntax and outputs of all statistical models. We used the lm() command from R base to build linear models, the lmer() command from the *lme4* package (73) for linear mixed-effect models, the nls() command from R base for non-linear regressions, and the glmer() command from the *lme4* package to build generalized linear mixed-effect models. We used the lsmeans() command from the *emmeans* package (74) for posthoc pairwise comparisons. To test the effect of the response variables, we either used the summary() command from R base or an ANOVA with the Anova() function of the *car* package (75). Whenever needed, we used a square root transformation to ensure that the residuals of the models followed a normal distribution. Supplementary Table 1 provides for each experiment the fixed and random variables that were included in the models. We included time as a fixed variable in the analyses of all the experiments that involved the collection of data at multiple time points. Whenever the analyses could not detect an interaction between time and the treatment of interest, we pooled the numbers of scans with brood care over the multiple time points. Whenever the analyses could detect an interaction between time and the treatment of interest, we investigated the effect of time separately for the different levels of the treatment variable (Supplementary Table 1).

### Detailed experimental descriptions

Below we provide experiment-specific information that deviates from the general procedures. Supplementary Table 2 provides an overview of the collection and keeping conditions of queens.

#### Brood production of founding queens

We used 24 founding queens collected in July 2019 on the campus of the Johannes Gutenberg University of Mainz, Germany (from now on referred to as JGU Mainz). We used a stereomicroscope to count for each queen the number of eggs, larvae, pupae, and workers three times per week for 93 days after the nuptial flight.

#### Effect of worker emergence

We used 12 founding queens collected in July 2017 on the campus of the JGU Mainz. We kept the queens in their collection boxes with humid paper towel for seven days before transferring them into individual glass tubes. The queens were filmed inside their tubes with all their brood for 15 minutes before worker emergence (35 days after the nuptial flight), and after worker emergence (56 days after the nuptial flight).

#### Effect of experimental manipulations of worker presence and feeding state

We used 54 founding queens collected in June 2018 near the campus of the JGU Mainz. The queens were kept in glass tubes at 25°C and approximately 80% humidity with a 12h/12h light/dark cycle. Just before the start of the experiment, all pupae were removed to prevent the emergence of workers during the experiment (all eggs and larvae were left in the tubes). The experiment started 30 days after collection. The queens were divided into four treatments: (1) queens were fed and were given five workers (n=14), (2) queens were fed and received no workers (n=11), (3) queens were not fed and were given five workers (n=15), and (4) queens were not fed and received no workers (n=14). For the feeding treatment, queens were hand-fed for 5-10 minutes with artificial food with a 2:1 (carbohydrate:protein) ratio (70). We filmed the queens in their tubes for at least 30 minutes twice a day (morning and afternoon) over seven consecutive days (14 time points per queen).

#### Effect of worker removal on established queens that had workers for 3 days

We used 48 founding queens collected in July 2019 in three locations in and around Mainz (Mainz-Bretzenheim, Mainz-Marienborn and campus of the JGU Mainz). The experiment started 55 days after collection, and all pupae were removed once 37 days after collection to ensure that no workers emerged before the start of the experiment. We assigned the queens to three different treatments: (1) queens with five larvae and five added workers (n=16), (2) queens with five larvae and without workers (n=16) and (3) queens with five larvae that formerly had five workers (workers removed; n=15). In the “workers removed” treatment, queens had been given five workers in their glass tubes three days prior to the experimental setup, and all tubes were filmed just before the experimental setup. Then, the queens were filmed in the observation arenas 24 and 48 hours after experimental setup.

#### Effect of worker removal on established queens that had workers for 2 years and 6 months

We used 12 colonies that were established in the laboratory by founding queens collected in July 2017 on the campus of the JGU Mainz. The experiment started 932 days after collection. All colonies were treated the same way. First, we filmed each queen with five of its larvae and five of its workers to allow direct observations of the queen behavior and standardize worker number across colonies. Then, we removed the last five workers, and filmed the queens 24 and 48 hours after experimental setup.

#### Effect of worker removal on established queens that had workers for 3 years and 2 months

We used 24 colonies that were established in the laboratory by founding queens collected in July 2017 on the campus of the JGU Mainz. The experiment started 1162 days after collection. The experimental setup consisted of one queen, five of its larvae and five of its workers in observation arenas. Then, we gradually removed one worker per day over the course of six days, until reaching zero workers. We filmed the observation arenas every day, and removed the workers after each video recording.

#### Effect of workers in unmated queens

We collected 27 unmated queens from field colonies in June and July 2021 near the campus of the JGU Mainz, and kept them at 24°C and 80% humidity in darkness. The experiment started right after collection. Each unmated queen was observed with five foreign larvae and either five (n=14) or zero (n=13) workers. We filmed the unmated queens six and 24 hours after the experimental setup.

#### Effect of worker number

We used 76 founding queens collected in July 2019 on the JGU campus. The experiment started 35 days after collection. We set up one queen with five larvae and either zero (n=14), one (n=18), two (n=15), three (n=13), or five (n=16) workers. We recorded the queens 24, 48 and 72 hours after the start of the experiment. A similar experiment was conducted using 80 founding queens collected in July 2020 in several locations in Mainz (Marienborn, Mainz-Hechtsheim, and Mainz-Oberstadt). This second experiment started 47 days after collection. For the last experimental setups of this second experiment, all pupae were removed to ensure that no workers naturally emerged before the experiment was completed. The queens were provided with 15 larvae and either zero (n=16), one (n=16), two (n=16), three (n=16), or five (n=16) workers, and were also filmed 24, 48 and 72 hours after the experimental setup.

#### Effect of worker cuticular hydrocarbons (CHC)

We used 36 queens collected in July 2019 on the campus of the JGU Mainz. The experiment started 40 days after collection. We produced five CHC extracts, each from 100 workers collected in a field colony around Mainz, Germany. The workers were sedated with CO_2_, transferred into glass vials, and immersed in *n*-hexane 10 minutes while occasionally swaying the vials. The liquid was transferred to a micro insert and completely exhausted under a gentle nitrogen stream. The CHC were then re-dissolved in 50μl of hexane and stored at 4°C until used in the experiments. We applied either 10μl of the worker CHC extracts (treatment) or 10μl of pure *n*-hexane (control) on glass beads (∼1.5mm, Roth GmbH). Five glass beads were placed on a bowl made of aluminum foil (5mm diameter) to ensure that the CHC extracts would not soak into the plaster below. One queen and five larvae were placed in each observation arenas, together with either treatment (n=18) or control (n=18) beads. We filmed the observation arenas three, six and 24 hours after experimental setup.

#### Effect of former presence of workers

We used 36 founding queens collected in July 2019 on the campus of the JGU Mainz. The experiment started 42 days after collection. Each queen was placed together with five larvae in an observation arena that used to contain 20 field-collected workers for 48 hours (workers removed just before the start of the experiment, n=18) or in a clean observation arena (n=18). We filmed the observation arenas six and 24 hours after the experimental setup.

#### Effect of the presence of dead workers

We used 54 founding queens collected in July 2020 in Ingelheim and Mainz, Germany. The experiment started 65 days after collection. The tubes containing the queens were regularly checked prior to the experiment to remove any pupae, thus ensuring that no workers had emerged before the experimental setup. We assigned the queens to four treatments: one queen with five larvae and (1) five dead workers (n=18), (2) five living workers (n=18) and (3) without any workers (n=18). Dead workers were obtained by placing 90 workers at -80°C for two hours, and then at -20°C for 24 hours before the experimental setup. Freshly killed, frozen ants show a similar CHC profile as living ants^28-30^. We recorded the observation arenas two, six and 24 hours after experimental setup.

#### Effect of workers separated by a wire mesh

We used 43 founding queens collected in July 2020 in Ingelheim and Mainz, Germany. The experiment started 52 days after collection. The tubes containing the queens were regularly checked prior to the experiment to remove any pupae, thus ensuring that no workers had emerged before the experimental setup. In this experimental setup, we modified the observation arenas by adding a wire mesh cylinder (0.2mm thick, 0.2mm mesh size, 2cm cylinder diameter) in the middle of the arena (Supplementary Figure 11). The wire mesh enabled antennation between the queen and the separated workers, but the workers had no close physical interactions with neither the queen nor the larvae. We assigned the queens to the following treatments: one queen and five larvae were placed outside the wire circle and the three workers were placed (1) on the inside of the circle, thus separated from the queen and brood (workers separated; n=13), or (2) on the outside of the circle together with the queen and brood (workers present; n=14) or (3) without any workers (workers absent; n=16). We filmed the observation arenas two, six and 24 hours after experimental setup.

#### Effect of methoprene and precocene I and II

We used 160 queens collected in July 2020 in Ingelheim, Mainz-Bretzenheim and the city center of Mainz, Germany. The experiment started 52 days after collection for the methoprene experiment, and 66 days after collection for the precocene I and II experiment. For the methoprene experiment, queens were treated with (1) methoprene at 1.5μg/μl (PESTANAL®, a commonly used JH analog(76); n=35) or (2) acetone (solvent; n=35). For the precocene I and II experiment, queens were treated with (1) precocene I at 1.5μg/μl (Sigma; JH inhibitor; n=30), (2) precocene II at 1.5μg/μl (Cayman; JH inhibitor; n=30) or (3) acetone (solvent; n=30). To treat the queens, we attached them to an eraser with a fishing line, and applied 1μl of the treatment on their thorax with a glass pipette. Each queen was treated every morning for four consecutive days. In the first two days, we recorded the observation arenas without workers. On the third day, before the application of the treatments, we added five callow workers to half of the queens, and kept them with these workers for another two days. The observation arenas were recorded two and six hours after each treatment. Note that the precocene II results are reported in Supplementary Figure 8 and Supplementary Table 1.

#### Effect of worker removal on established *Temnothorax nylanderi* queens

We collected 21 *T. nylanderi* colonies in April 2021 in the Lenneberg forest of Mainz, Germany. In the laboratory, each colony was transferred into plastered nest boxes containing an artificial nest consisting of a Plexiglas perimeter (3mm high) with an entrance, sandwiched between two microscope slides (7.5cm × 2.5cm × 0.5cm). The colonies were kept according to the general procedures, and were fed twice a week with a drop of honey, half a cricket and water. The experiment started two weeks after collection. The treatments consisted of queens with five larvae and (1) five of their own workers (n=11) and (2) no workers (n=10). We used CO_2_ to anesthetize and transferred the ants into the observation arenas. We filmed the observation arenas every 24 hours for five consecutive days.

#### Effect of larvae on egg production in founding and established queens

We used 66 *L. niger* queens collected in July 2019 on the campus of the JGU Mainz. The queens were distributed into the following treatments: (1) founding queens with larvae (n=18), (2) founding queens without larvae (n=18), (3) established queens with larvae (n=15), and (4) established queens without larvae (n=15). Founding queens were queens that had not yet produced pupae or workers, and were tested eight days after the nuptial flight. Established queens were queens that had produced at least five workers, and were tested 82 days after the nuptial flight. To test the effect of larvae, we used five larvae collected in field colonies near the campus of the JGU Mainz. We recorded the number of eggs once a day for five consecutive days.

## Acknowledgments

We thank former and current members of the Behavioural Ecology and Social Evolution group at the JGU Mainz for help with the collection of founding queens, and fruitful discussions on the project.

## Author contributions

Conceptualization: RL

Methodology: RL

Investigation: VM, MNP, MC, EF, EMT, FR, JG, LM, MR, DU, RL

Analysis: VM, MNP, MC, RL

Funding acquisition: RL

Project administration: RL

Supervision: RL

Writing - original draft: VM, MNP, MC, RL

Writing - review and editing: VM, MNP, MC, RL

## Data and materials availability

All data are available in the main text or the supplementary materials.

## Supplementary materials

Supplementary Figures (S1 to S11)

Supplementary Tables (S1 to S2)

