## Supplementary Figures for "Ontogeny of superorganisms: Social control of queen specialization in ants"

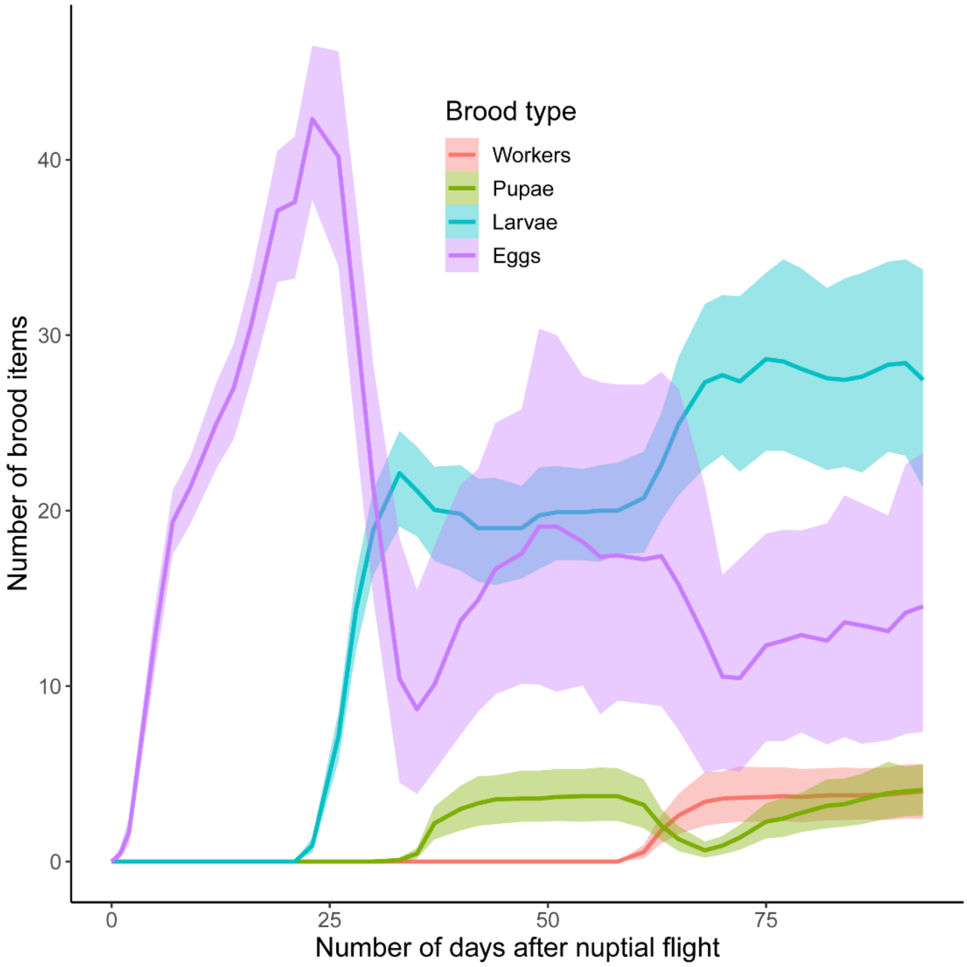


| | Eggs | Larvae | Pupae | Workers |
|---|---|---|---|---|
| % of surviving queens | 91.67% (22/24) | | | |
| % of queens that produced each brood item | 100% (22/22) | 95.45% (21/22) | 77.28% (17/22) | 77.28% (17/22) |
| mean±sd day of first occurrence | 2.91±1.82 | 24.43±1.53 | 36.53±2.58 | 62.88±2.52 |
| mean±sd maximum number of each brood item produced | 49.73±17.07 | 33.95±13.57 | 6.53±3.68 | 5.26±3.95 |

**Supplementary Figure 1.** Brood production of 22 founding queens. The eggs, larvae, pupae and workers of each queen were counted in their glass tubes three times per week for 93 days after the nuptial flight. Additional information on queen survival and brood production can be found in the table below the figure.

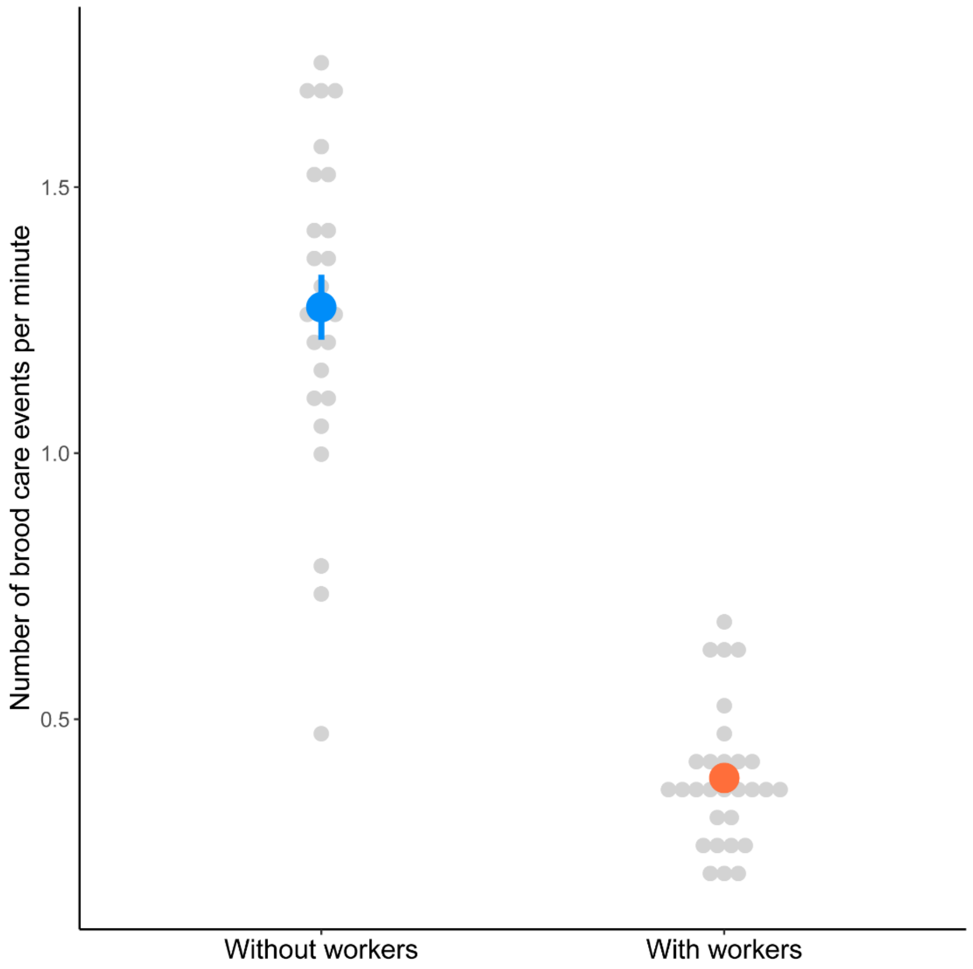


**Supplementary Figure 2.** Effect of workers on the brood care behavior of founding queens 20 hours after the experimental manipulation. This corresponds to the first time point of the experiment "Effect of experimental manipulations of worker presence and feeding status" (Supplementary Table 1). The experimental addition of workers already reduced the brood care behavior of founding queens 20h after the experimental manipulation (ANOVA: $F=221.28$, $P<0.0001$).

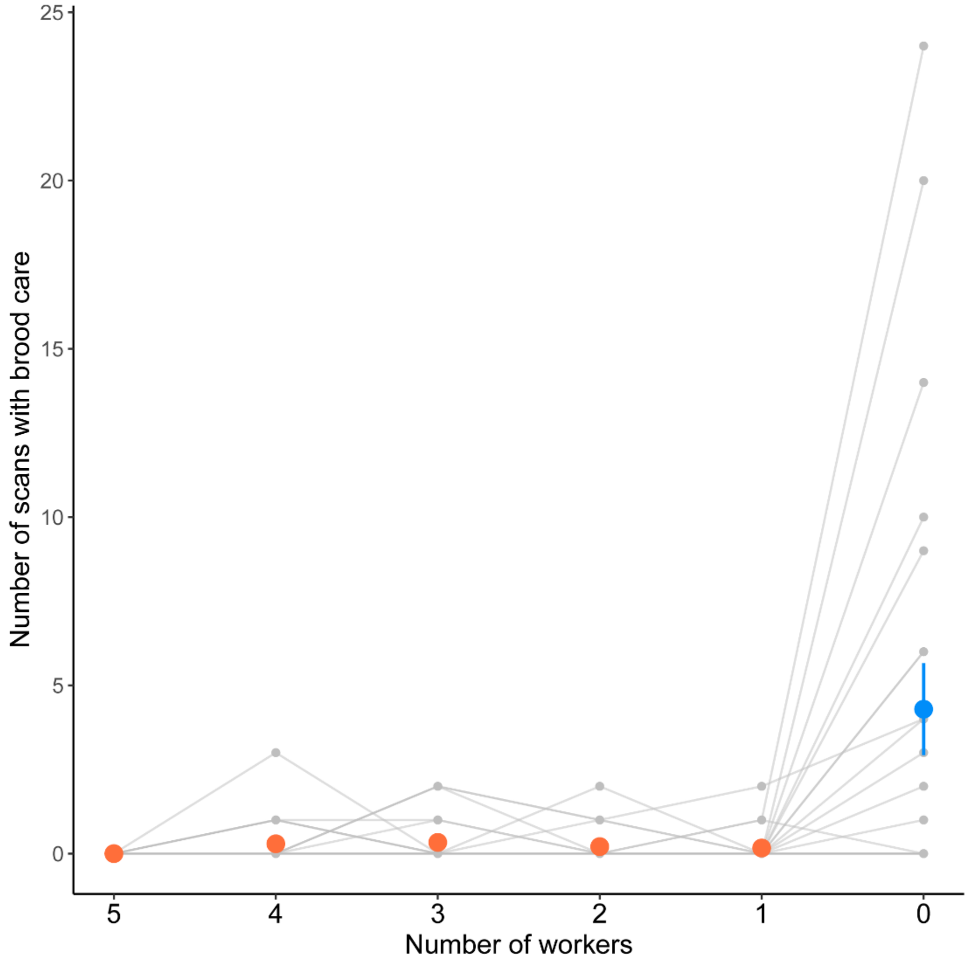


**Supplementary Figure 3.** Brood care behavior of established queens (three years and two months after their nuptial flight) in response to the sequential removal of their workers over the course of six days. The experimental setup consisted of one queen, five of her larvae and five of her workers in observation arenas. We gradually removed one worker per day over the course of six days, until no workers remained (the workers were removed after each daily recording). We built a linear mixed-effect model with the number of workers as fixed factor, and queen identity and batch as random factors (Supplementary Table 1). Since the number of workers influenced the brood care behavior (ANOVA: $\chi^2$=55.53, P<0.0001), we further investigated the number of workers needed to produce the effect. We calculated all posthoc pairwise comparisons, and found that all categories differed from the category without workers (all t>5.2; all P<0.0001).

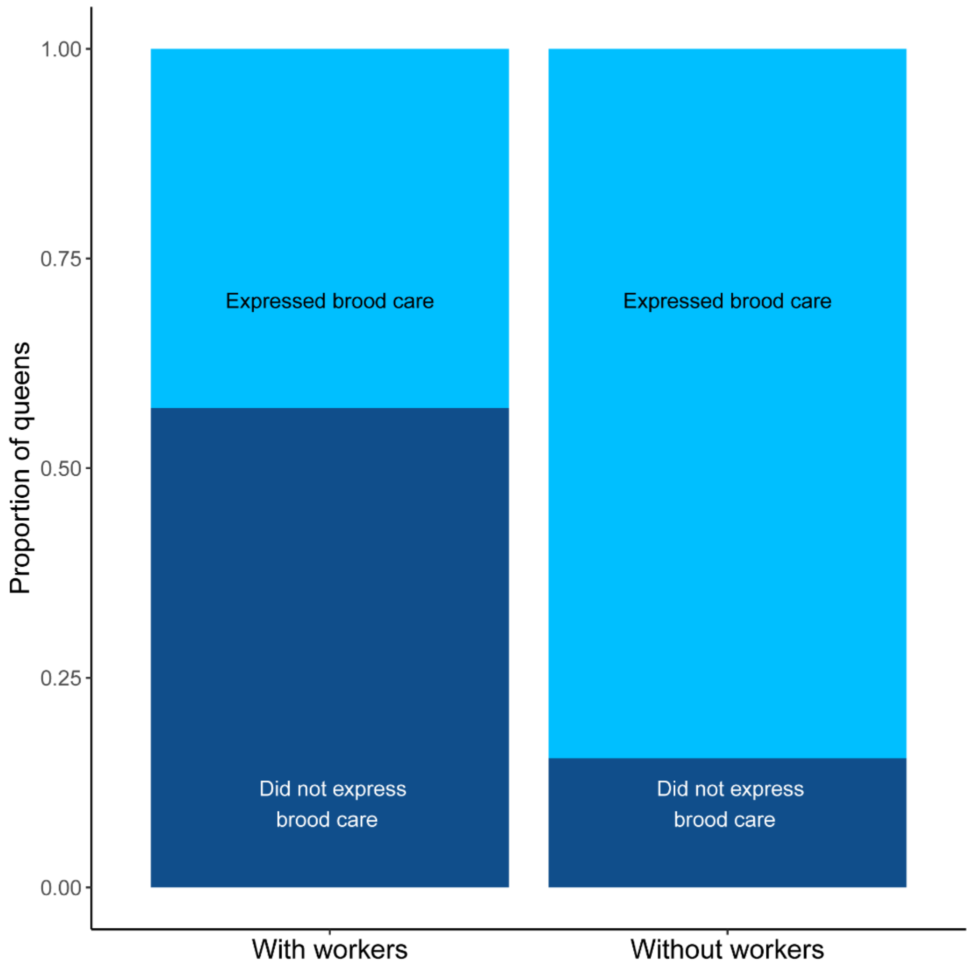


**Supplementary Figure 4.** Proportion of unmated queens expressing brood care behavior in the presence or absence of workers. Each of the 27 unmated queens was observed with five foreign larvae and with either five workers (n=14) or zero workers (n=13). We filmed the observation arenas six and 24 hours after experimental setup. Due to the low number of brood care events overall, we pooled the two time points, transformed the brood care events into binary data, and created a generalized linear mixed-effect model (family: binomial) with worker presence as fixed factor and batch as random factor (Supplementary Table 1). We found moderate evidence for the proportion of unmated queens expressing brood care to be lower for unmated queens kept with workers compared to unmated queens kept without workers (ANOVA: $\chi^2$=4.49, P=0.034).

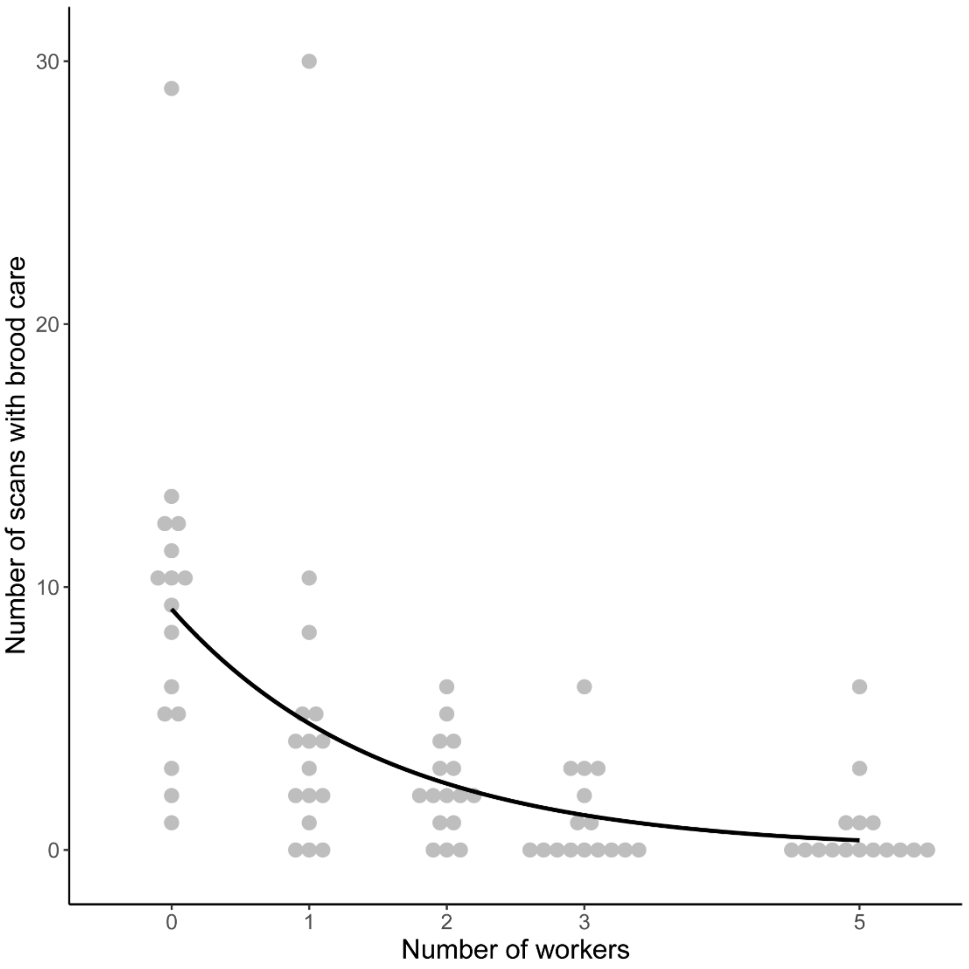


**Supplementary Figure 5.** Workers induce queen specialization in a dose-dependent manner. Eighty founding queens were provided with zero (n=16), one (n=16), two (n=16), three (n=16) or five (n=16) workers, as well as 15 larvae, and were filmed 24, 48 and 72 hours after experimental setup. We built a linear mixed-effect model with time after setup, number of workers and their interaction as fixed factors, and queen identity and tray as random factors. Since we did not detect any interaction between time and treatment, we pooled the data from the three time points, and fitted a non-linear model with the function:

$$Brood\ care = a * e^{b * number\ of\ workers}$$

We found the parameter "b" to be negative and different from 0, thus revealing a negative correlation between the number of workers and the level of brood care expressed by queens, both when queens without workers were included (t=4.19, P<0.0001) and excluded from the analysis (t=2.37, P=0.021).

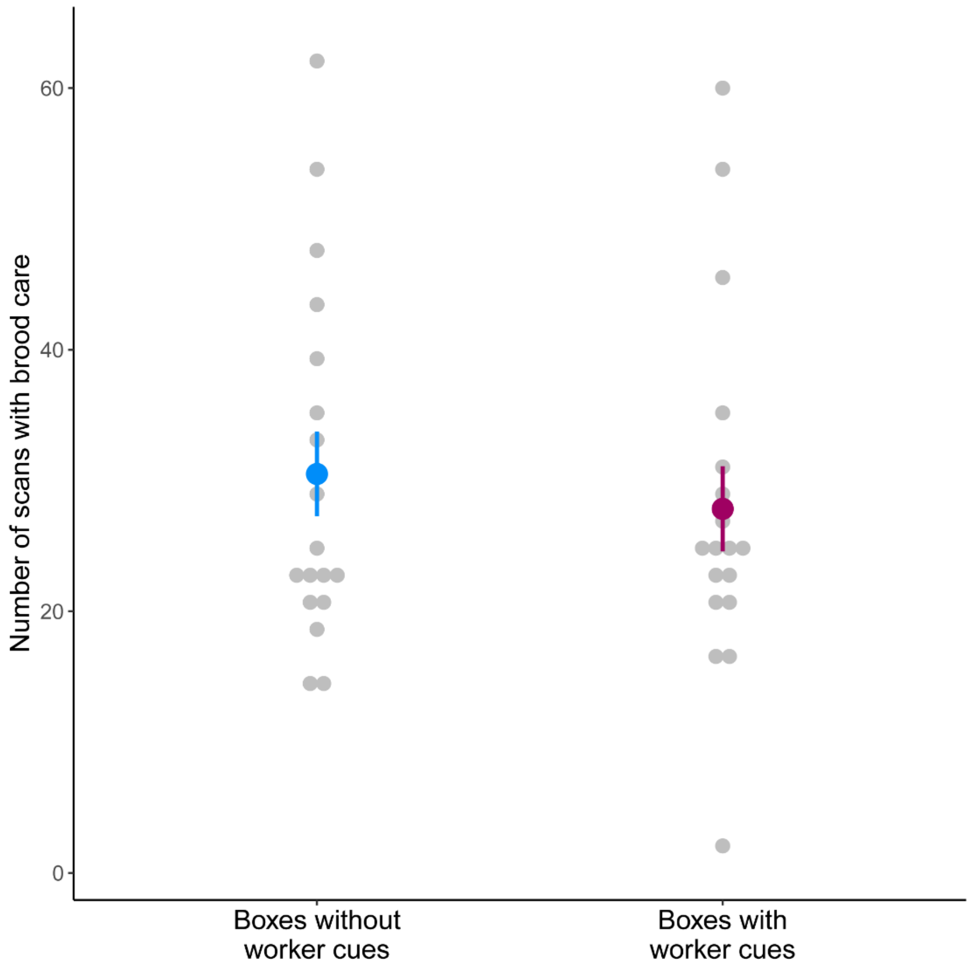


**Supplementary Figure 6.** No effect of former presence of workers on the brood care behavior of founding queens. Thirty-six founding queens were assigned to one of the following treatments: one queen and five larvae were placed in an observation arena that used to contain 20 field-collected workers for 48 hours (the workers were removed just before the start of the experiment, n=18) or kept in clean observation arenas (n=18). The queens were recorded six and 24 hours after experimental setup. Since we found no interaction between time and treatment, the two time points were pooled. We built a linear mixed-effect model using treatment as fixed factor and the tray number as random factor. We could not detect any effect of the presence of worker cues (ANOVA: $\chi^2$=0.48, P=0.49).

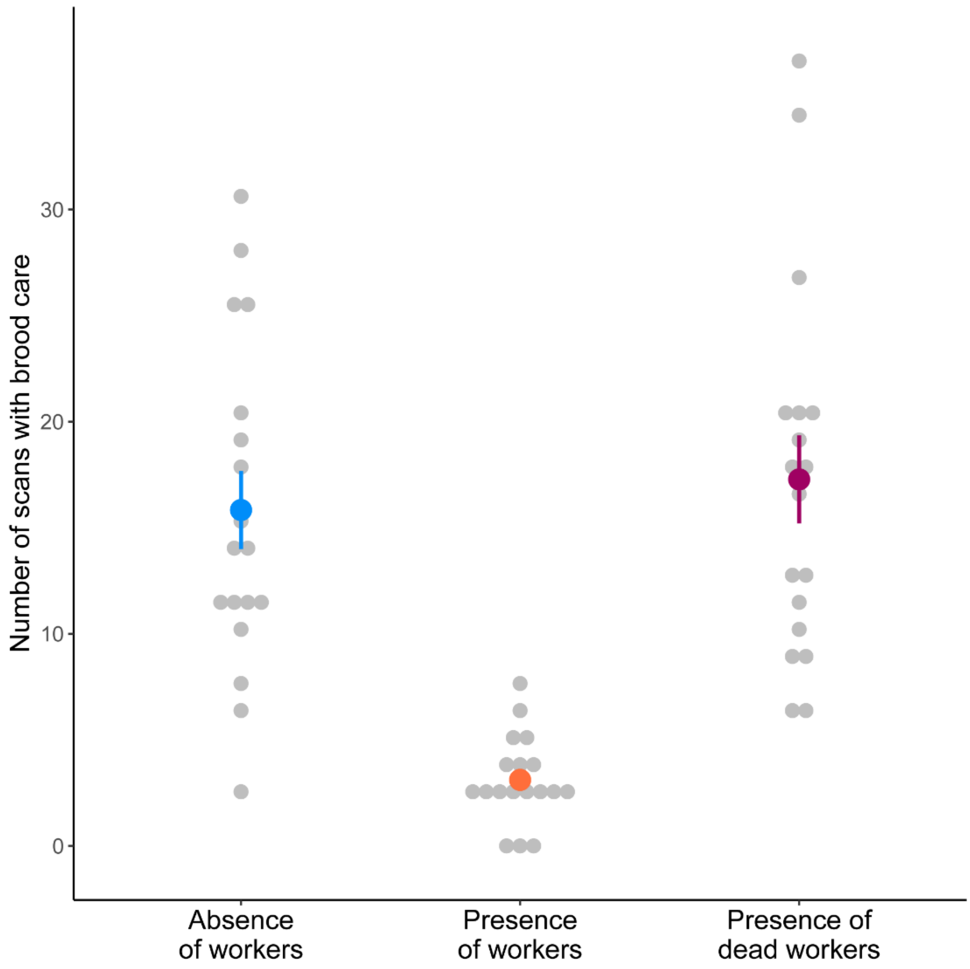


**Supplementary Figure 7.** Effect of dead worker presence on brood care of founding queens. We assigned 54 founding queens to four treatments: one queen with five larvae and (1) five dead workers (n=18), (2) five living workers (n=18) and (3) without any workers (n=18). The dead workers used in the experiment were obtained by placing 90 workers at -80°C for two hours, and then at -20°C for 24 hours before the experimental setup. We recorded the observation arenas two, six and 24 hours after experimental setup. We built a linear mixed-effect model with time after experimental setup and treatment as fixed factors, and the queen identity and tray as random factors (Supplementary Table 1). Since we could not detect an interaction between time and treatment, we pooled the three time points, and included in the model treatment as fixed factor and tray as a random factor. We found a main effect of treatment (ANOVA: $\chi^2$=73.85, P<0.0001). Founding queens in presence of living workers exhibited less brood care compared to both queens without workers (t=7.14, P<0.0001) and with dead workers (t=7.71, P<0.0001), while there was no difference in brood care between queens without workers and with dead workers (t=0.57, P=0.84).

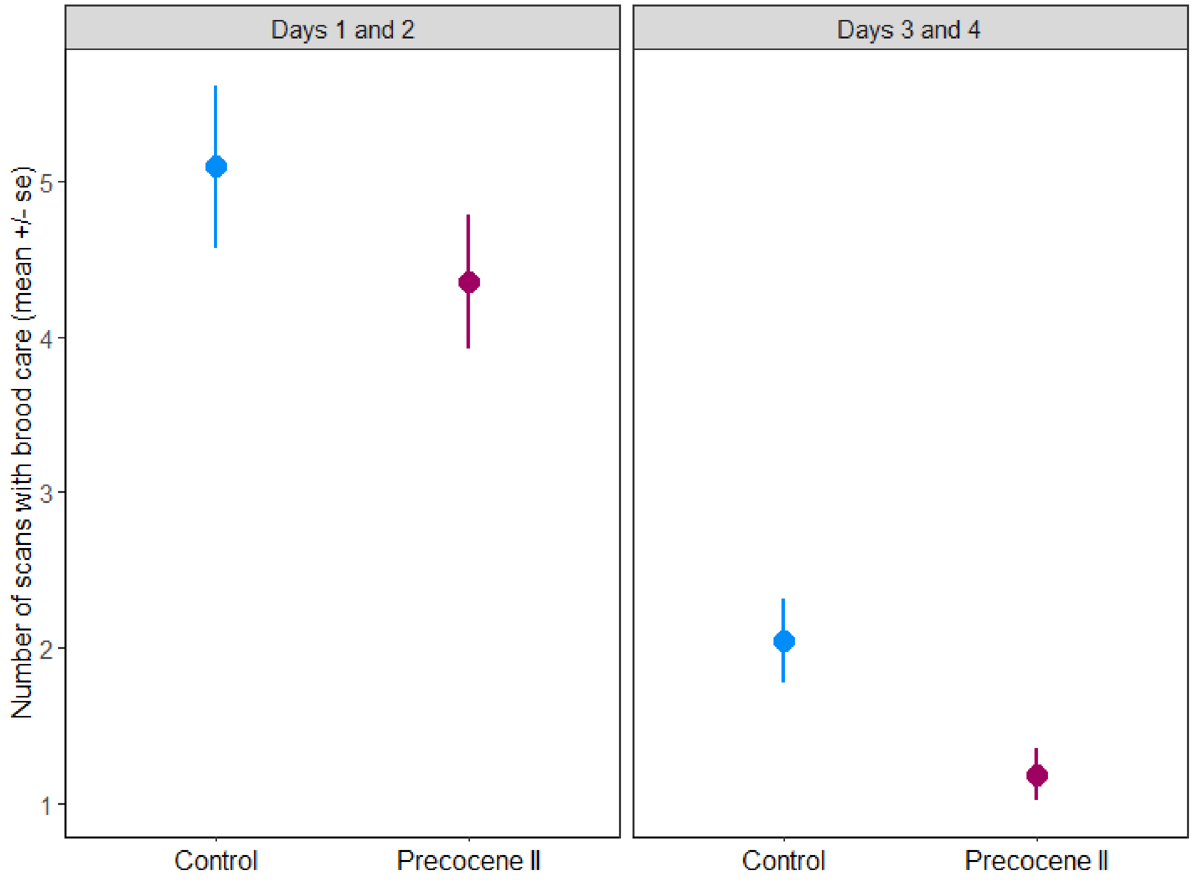


**Supplementary Figure 8.** Effect of precocene II on brood care behavior in founding queens. We used 90 founding queens that were treated with either precocene II at 1.5µg/µl (Cayman, n=30) or the solvent as control (acetone, n=30). To apply the treatments, we attached the queens to an eraser with a fishing line, and applied 1µl of the respective treatment on their thorax with a glass pipette. The treatments were applied daily in the morning for four consecutive days. In the first two days, we recorded the queens with larvae but without workers. On the third day, before applying the treatments, we added five callow workers to half of the queens, and kept them with these workers for another two days. The queens were recorded two and six hours after each treatment. All the statistical models and outputs are available in Supplementary Table 1. We built separate linear mixed-effect models (one for days 1 and 2, another for days 3 and 4) using time, time since first treatment and treatment as fixed factors (and all interactions) as fixed effects, and queen identity, tray and batch as random factors. In addition, for days 3 and 4, we added workers presence as another fixed factor. In the first period (days 1 and 2), we did not detect any effect of the fixed factors (Supplementary Table 1). In the second period (days 3 and 4), we found moderate evidence for an effect of the precocene II treatment (ANOVA: $\chi^2$=5.82, P=0.016).

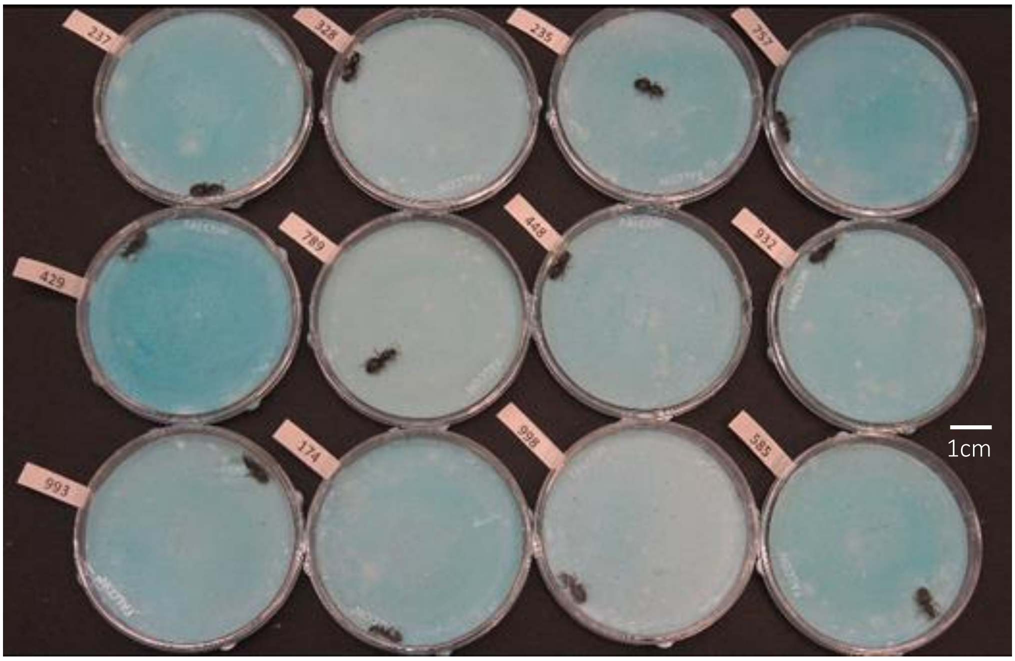


**Supplementary Figure 9.** One experimental tray with 12 observation arenas, which were recorded simultaneously with a single camera. Queens were assigned a random number to allow for blind observations, and we ensured that all treatments were equally represented and evenly distributed across and within trays.

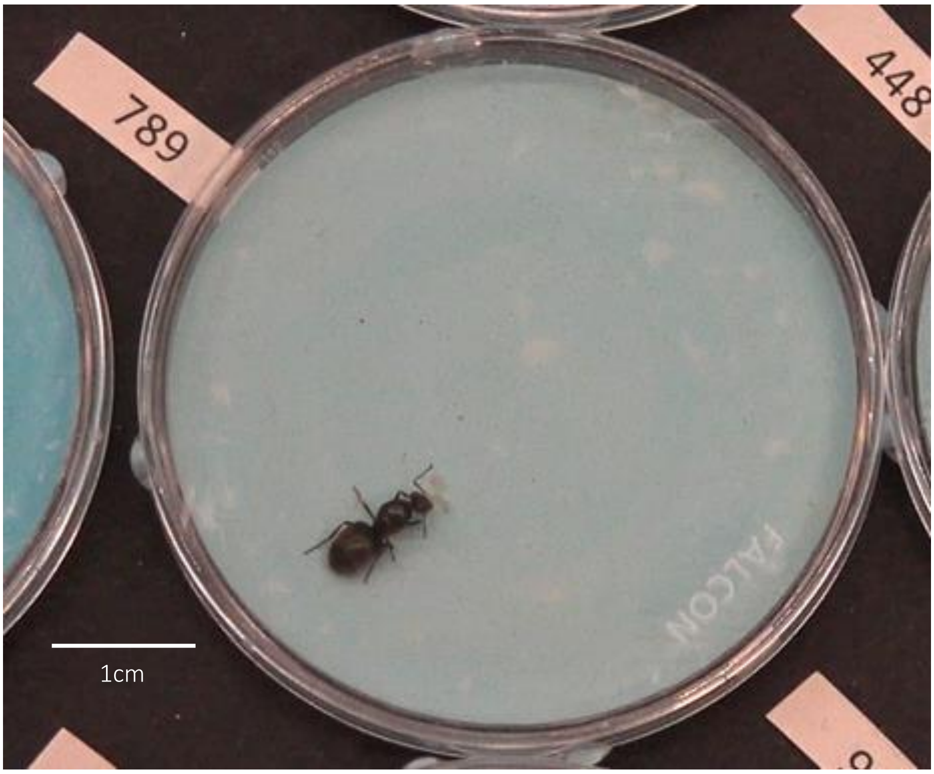


**Supplementary Figure 10.** Resolution of the video when zooming in on one observation arena with one queen and five own larvae. The queen brood care behavior was assessed at this zoom level. We used blue-colored plaster to improve contrast and visibility of the larvae.

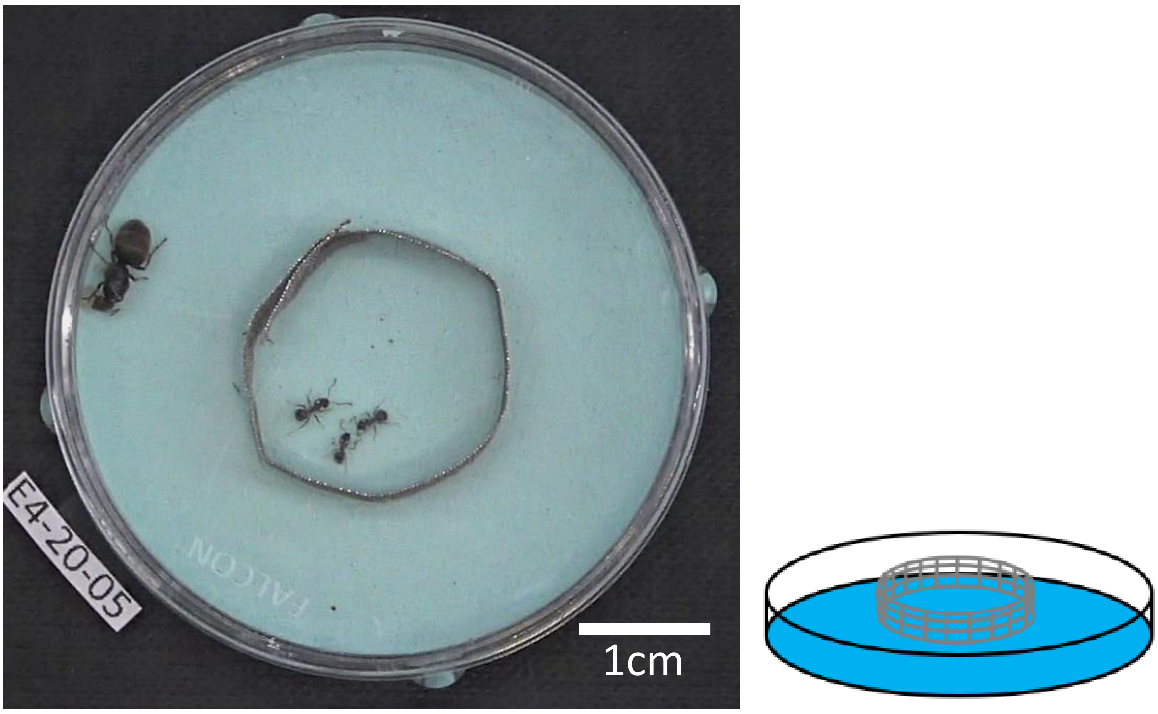


**Supplementary Figure 11.** Left: Experimental setup to test workers separated from the queens with a circular wire mesh (wire mesh cylinder: 0.2mm thick, 0.2mm mesh size, 2cm cylinder diameter**). The queen brood care** behavior was scored at this zoom level. Right: Schematic side-view of the observation arena with wire mesh. We used blue-colored plaster to improve contrast and visibility of the larvae.
